# Auditory aversion in absolute pitch possessors

**DOI:** 10.1101/2020.06.13.145029

**Authors:** Lars Rogenmoser, H. Charles Li, Lutz Jäncke, Gottfried Schlaug

**Author notes:** Address correspondence to Lars Rogenmoser, Department of Neuroscience, Georgetown University Medical Center, 3970 Reservoir Road NW, Washington, DC 20007, USA.

## Abstract

Absolute pitch (AP) refers to the ability of identifying the pitch of a given tone without reliance on any reference pitch. The downside of possessing AP may be the experience of disturbance when exposed to out-of-tune tones. Here, we investigated this so-far unexplored phenomenon in AP, which we refer to as auditory aversion. Electroencephalography (EEG) was recorded in a sample of AP possessors and matched control musicians without AP while letting them perform a task underlying a so-called affective priming paradigm: Participants judged valenced pictures preceded by musical primes as quickly and accurately as possible. The primes were bimodal, presented as tones in combination with visual notations that either matched or mismatched the actually presented tone. Regardless of the prime condition, AP possessors performed more poorly in judging pleasant pictures and their EEG revealed later peaks at around 200 ms (P200) after prime onset. Their performance dropped when responding to pleasant pictures preceded by incongruent primes, especially when mistuned by one semitone. This interference was also reflected in an EEG deflection at around 400 ms (N400) after picture onset, preceding the behavior responses. These findings suggest that AP possessors process mistuned musical stimuli and pleasant pictures as affectively unrelated with each other, supporting an aversion towards out-of-tune tones in AP possessors. The longer prime-related P200 latencies exhibited by AP possessors suggest a delay in integrating musical stimuli, underlying an altered affinity towards pitch-label associations.

## 1. Introduction

Absolute pitch (AP), the ability to identify the chroma of a tone without the aid of any reference (Takeuchi and Hulse 1993), is sparsely distributed in the population (approximate <1%) but yet bears phylogenetic and ontogenetic significance. Songbirds fail to demonstrate octave generalization while experiments confirm this skill in rhesus monkeys (Cynx 1993; Wright et al. 2000), suggesting that hearing with a preference for absolute over relational cues is conserved in humans (Levitin and Rogers 2005). Humans prefer absolute auditory cues during early infancy, shifting towards a preference for relational ones as they mature (Saffran and Griepentrog 2001). A genetic component in combination with specific learning factors such as early music engagement and exposure to language, and possibly the presence of a particular brain anatomy prevent the maturation-related decline in the ability of processing pitches absolutely and thus account for AP acquisition (Gregersen PK, Kowalsky E, Kohn N 1999; Gregersen et al. 2001; Deutsch et al. 2006). Similar to the successful language acquisition that requires early language exposure during a child’s development (Newport 1990), AP may only emerge on condition that music engagement takes place during a critical period, a maturational stage at which cognitive processing still functions in a rather unidimensional mode and the brain is still highly malleable and responsive to environmental inputs (Chin 2003; Russo et al. 2003). As a result, adult AP possessors show an increased left-sided asymmetry of the planum temporale (Schlaug et al. 1995; Keenan et al. 2001; Wilson et al. 2009); the structural asymmetry might create a functional dominance directing auditory stimuli to categorization centers within the temporal lobe such as the superior temporal sulcus (Loui et al. 2010; Jäncke, Langer, et al. 2012). A further brain structure related to AP is the posterior dorsal frontal cortex. This brain region might potentially drive conditional associative memory and, in the context of AP, be responsible for the process underlying the association between categorized pitches and verbal labels or other sensorimotor codes (Zatorre and Beckett 1989; Zatorre et al. 1998). In AP possessors, within this left-sided frontotemporal network an interplay is detectable even at rest in electroencephalography (EEG) recordings (Elmer et al. 2015), suggesting functional optimization that may enable the automatic nature of AP. Automaticity is, however, accompanied by the difficulty in suppression, as many studies using interference tasks have demonstrated, revealing a drop in identification performance under incongruent trial conditions among AP possessors (Itoh et al. 2005; Akiva-Kabiri and Henik 2012; Schulze et al. 2013; Ziv and Radin 2014; Leipold, Greber, et al. 2019). Further studies revealed that in comparison to non-AP (NAP) subjects, AP possessors are disadvantaged in recognizing transposed melodies or in identifying musical intervals within out-of-tune contexts (Miyazaki 1993; Miyazaki and Rakowski 2002). AP possessors require more cognitive load for processing out-of-tune tones, this certain vulnerability towards mistuned tones increasing the later the onset of musical training took place during childhood (Rogenmoser et al. 2015). From an affective perspective, this vulnerability is corroborated by anecdotal evidence suggesting that AP possessors become agitated when exposed to out-of-tune tones (Levitin and Rogers 2005). Here, we investigated this so far unexplored phenomenon in AP, which we refer to as auditory aversion. Understanding the mechanisms underlying auditory aversion might advance research on cognitive-affective development. It is further crucial given that some AP possessors report a distortion in frequency perception as a consequence of aging (Vernon 1977; Athos et al. 2007). Thus, aversion would no longer exclusively be triggered externally by certain out-of-tune tones but additionally internally by perceiving in-tune tones as out-of-tune. In the present study, we measured the affective reactions in response to musical stimuli in a sample of AP possessors and in one of matched control participants by recording EEG during the performance of a task underlying an affective priming paradigm (Fazio 2001). The affective priming procedure involves the sequential presentation of two valenced stimuli, namely the preceding so-called primes and the following so-called targets. The task is to handle the targets whereby the preceding primes are either affectively related or affectively unrelated to them. It is well documented that affectively unrelated prime-target pairs induce interference that varies as a function of the extent of involved emotional experience (Fazio 2001). In EEG, this interference is reflected by a negative-going deflection around 400 ms after target-onset, which is referred to as the N400 (Kutas and Hillyard 1980; Zhang et al. 2010). EEG has an excellent time resolution and has been insightful in capturing the early perceptual and cognitive processes underlying AP (Elmer et al. 2013; Rogenmoser et al. 2015; Greber et al. 2018; Burkhard et al. 2019; Leipold, Oderbolz, et al. 2019). In the present study, the participants were instructed to judge as quickly and accurately as possible whether pictures (targets) were either pleasant or unpleasant by pressing one of two response buttons. In the experimental condition, these targets were preceded by musical stimuli (primes) which were bimodal, comprising visually presented notations in combination with auditorily presented tones. The auditory counterpart either matched the notations (congruent) or deviated from them (incongruent) in a higher semitone (sharp) or quarter-tone. Based on the assumption that out-of-tune tones trigger aversion in AP possessors, we predicted an interference in the AP possessors’ response behavior accompanied by a greater N400 deflection in conditions in which the incongruent primes preceded pleasant targets.

## 2. Materials and methods

### 2.1 Participants

Twenty-one AP possessors (14 males) and 21 matched NAP control musicians (13 males) participated in this study. The two samples were comparable regarding age (*t*_40_ = 0.80, *P* = 0.428, *d* = 0.25), general cognitive capability (*t*_39.91_ = 0.677, *P* = 0.502, *d* = 0.21) as measured by the Shipley–Hartford Retreat Scale (Shipley 1940), and comparable regarding the distribution of handedness (one left-hander in each sample) and of the sexes (*Χ^2^*_1_ = 0.10, *P* = 0.747). Both AP and NAP subjects commenced their musical training at a comparable age range (*t*_40_ = 1.64, *P* = 0.106, *d* = 0.51) and trained for a comparable number of years (*t*_40_ = 0.11, *P* = 0.910, *d* = 0.04). Across both samples, the subjects were comparably skilled in music performance, as confirmed using pitch and rhythm discrimination tasks underlying psychophysical adaptive staircase procedures (Loui et al. 2009; Fujii and Schlaug 2013). Regarding pitch discrimination, the minimum detectable frequency difference around the center frequency of 500 Hz was measured. The two samples performed at comparable pitch discrimination thresholds (*t*_40_ = 1.18, *P* = 0.244, *d* = 0.36). Regarding rhythm discrimination (i.e., Beat Interval Test (Fujii and Schlaug 2013)), sequences of 21 woodblock tones with varying interval lengths were presented. The sensory threshold for discriminating temporal changes as well as the performance in tapping in synchrony while adapting to temporal changes was measured. The two groups performed at comparable rhythm discrimination thresholds (*t*_31.83_ = 0.011, *P* = 0.992, *d* = 0.00) and with comparable tapping synchronization abilities (*t*_40_= 0.67, *P* = 0.507, *d* = 0.21). The values on characteristics and musical background are reported in Table 1. All subjects gave written informed consent, and the study was approved by the institutional review board of Beth Israel Deaconess Medical Center.

**Table 1.** Characteristics and data on the musical background of both samples.

|  | AP | NAP |
| --- | --- | --- |
| Age (years) | 22.38 (4.86) | 23.47 (3.94) |
| Cognitive capability (IQ scores) | 115.24 (8.62) | 113.47 (8.22) |
| Age at commencement of musical practice (years) | 6.14 (2.44) | 7.43 (2.60) |
| Duration of musical training (years) | 16.24 (5.70) | 16.05 (5.08) |
| Pitch discrimination threshold (Hz) at 500 Hz | 4.89 (5.43) | 7.55 (8.77) |
| Rhythm; perceptual discrimination threshold (ms) | 1.53 (1.72) | 1.54 (3.01) |
| Rhythm; tapping synchronization threshold (ms) | 1.61 (3.56) | 2.39 (4.03) |

Listed are the means with the standard deviations in brackets. All independent-samples $t$ -test calculated for each variable revealed values of $P > 0.1$ . AP: Absolute pitch, NAP: non-absolute pitch

### 2.2 Absolute pitch (AP) verification

AP was confirmed using an established pitch-labeling test (Keenan et al. 2001) in which 52 trials were presented. Each trial contained one computer-generated sine wave tone (500-ms duration with a 50-ms rise and decay time) with a fundamental frequency covering one octave from 370 Hz (F#3) to 739.97 Hz (F#4) in the equal-tempered Western scale. The participant’s task was to label each pitch by writing down the letter name on an answer sheet upon hearing each tone. The inter-tone interval was 2 s. The AP possessors performed (% correct responses) considerably better on this test (mean correct: 94.15, *SD* = 10.05) than the NAP subjects (mean correct 5.77, *SD* = 6.26; *t*_40_ = 34.21, *P* = 3.08×10^−31^, *d* = 10.56). NAP participants did not perform better than chance level (8.33%; *t*_20_ = –1.89, *P* = 0.075).

### 2.3 Experimental task and stimulus material

To measure the participants’ affective reaction in response to musical stimuli, EEG was recorded during performance of a task underlying an affective priming paradigm. In this task, the participants were instructed to judge as quickly and accurately as possible whether pictures (targets) preceded by musical stimuli (primes) were either pleasant or unpleasant by pressing one of two response buttons. The targets were 144 pictures (see Appendix) taken from the International Affective Picture System (Lang et al. 2008) on the basis of normative ratings (9-point scales) regarding the two affective dimensions of valence (1 = most unpleasant; 9 = most pleasant) and arousal (1 = most calm; 9 = most aroused). Half of the picture collection was pleasant (*M* = 7.60, *SD* = 0.41) while the other was unpleasant (*M* = 2.66, *SD* = 0.79; *t*_107.21_ = 46.91, *P* = 2.76×10^−73^, *d* = 7.82), but across both valences the pictures were controlled for arousal (unpleasant pictures: *M* = 5.59, *SD* = 0.71; pleasant pictures: *M* = 5.41, *SD* = 0.95; *t*_131.31_= 1.33, *P* = 0.185, *d* = 0.21). The preceding primes were bimodal, comprising visually presented notations in combination with auditorily presented pure tones. The set of presented notations comprised the scale of 12 subsequent notes ranging from F#3 to F#4. The auditory counterpart either matched the notations or deviated from them in a higher semitone (sharp) or quarter-tone. The 144 targets were presented 4 times in total; once per each matching condition and additionally once as control condition without any prime. Since the tuning of the musical stimulus is expected to impact the affective reaction exceptionally in AP subjects, the particular prime-target pairs in this experimental set-up reflect either affectively unrelated or affectively related conditions. Predicated on this constellation, 6 particular prime-target pair conditions result (see Table 2), within which each of the 12 scaled primes were paired 4 times with different targets (72 times per valence). The prime-target pairs as well as the order of trials was pseudorandomized. Each trial began with a fixation cross for gaze stabilization that was presented for a duration of 500 ms. The following primes were presented with a duration of 500 ms and the targets with one of 800 ms while the prime-target interval was of 200 ms. The inter-trial interval was jittered between 1000 and 1500 ms. The procedure of one trial is illustrated in Figure 1. The auditory stimuli included 10 ms of fade-in/out phases and were delivered via Sennheiser HD 205 headphones at the sound pressure level of 80 dB. The visual material was shown in the center of a PC laptop monitor. Stimuli presentation and behavior recordings were controlled by the software PsyTask. Before starting the actual experiment, the subjects were given 5 practice trials to get familiar with the task.

**Table 2.** Prime-target pair conditions.

|  |
| --- |
| Affectively related: |
| 1. Congruent prime + pleasant target |
| 2. Incongruent prime (semitone) + unpleasant target |
| 3. Incongruent prime (quarter-tone) + unpleasant target |
| Affectively unrelated: |
| 4. Congruent prime + unpleasant target |
| 5. Incongruent prime (semitone) + pleasant target |
| 6. Incongruent prime (quarter-tone) + pleasant target |

**Fig. 1.**
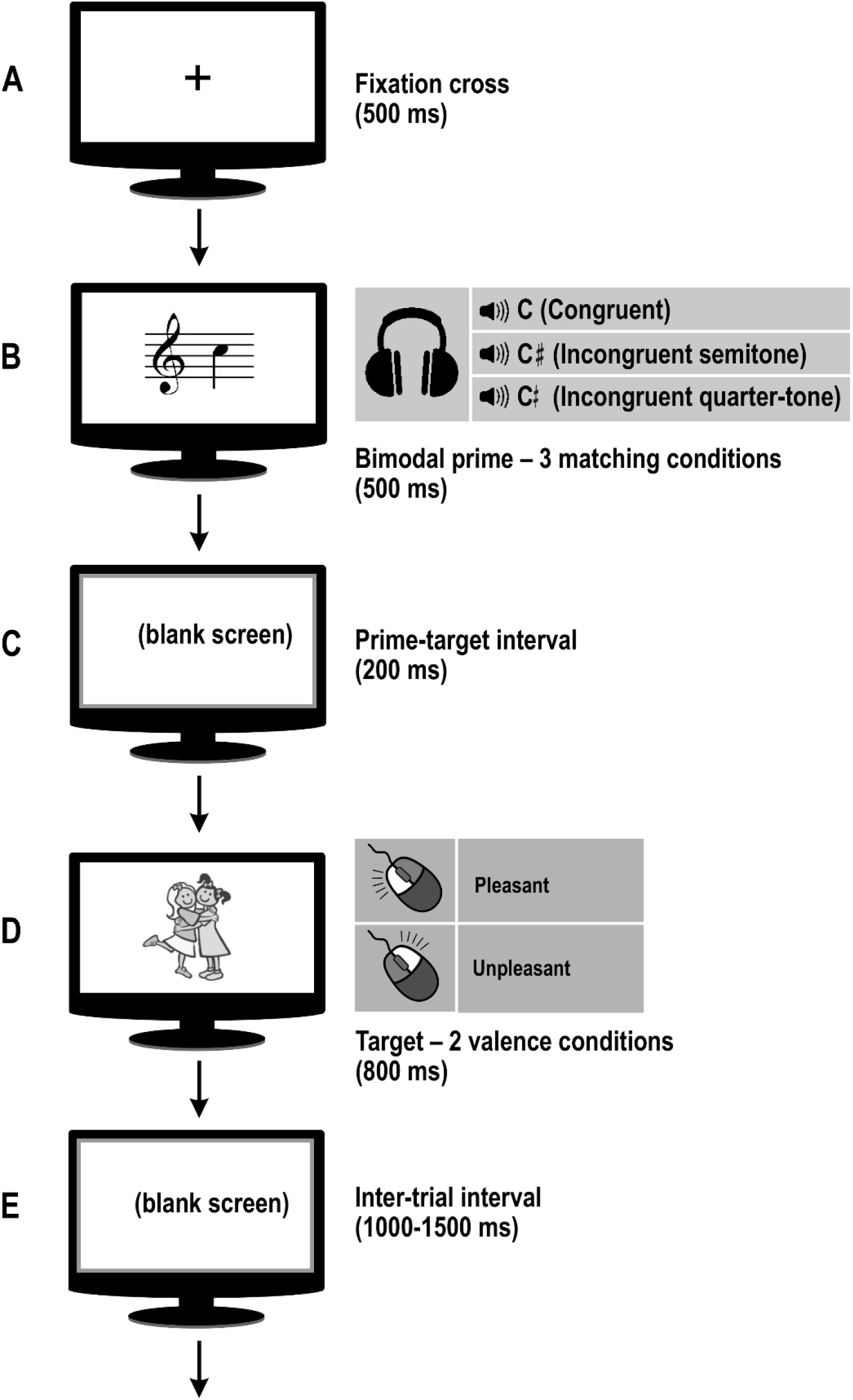
Schematic representation of the task. Each trial began with a fixation cross (A) that was presented on the monitor for 500 ms, followed by a prime (B) for a duration of 500 ms. The prime was bimodal, comprising visually presented notations in combination with auditorily presented pure tones via earphones. The auditory counterpart either matched the notations (congruent) or deviated from them (incongruent) in a higher semitone or quarter-tone. Afterwards, the monitor turned blank for 200 ms (C). Then, the target (D) followed, presented for 800 ms. The target was either a pleasant or an unpleasant picture taken from the International Affective Picture System. The participants were instructed to indicate as quickly and accurately as possible the valence of the target by pressing on either the right (pleasant) or left (unpleasant) mouse button. The next trial followed after the monitor turned blank for a duration jittered between 1000 and 1500 ms (E).

### 2.4 EEG recording

EEG was recorded via 19 channels (Ag/AgCl electrodes, BEE Medic GmbH) placed according to the “10-20” system with the following electrode locations: Fp1, Fp2, F7, F3, Fz, F4, F8, T3, C3, Cz, C4, T4, T5, P3, Pz, P4, T6, O1, O2. A further Ag/AgCl electrode embedded in the cap horizontally centered between Cz and Fz served as ground and two Ag/AgCl electrodes clipped on both earlobes as reference. EEG recording was done with the EEG NeuroAmpx23 amplifier and the ERPrec software. The signal was recorded with a sampling rate of 500 Hz and a bandpass filter of 0.1-100 Hz. Impedances were kept below 5 kΩ using conductive gel (Elecro-Gel).

### 2.5 Data analyses

#### 2.5.1 Preprocessing

Raw EEG data were imported into EEGLAB v.13.2.1 (Delorme and Makeig 2004), an open source toolbox running under MATLAB. The data were down-sampled to 250 Hz, band-pass filtered at 1-20 Hz and re-referenced to the mean of the T3 and T4 electrodes. Epochs were created ranging from −200 to 500 ms after prime-onset and from −200 to 1000 ms after target-onset, respectively. A baseline correction relative to the −200 to 0 ms pre-stimulus time period was applied. Epochs contaminated by unsystematic artifacts were rejected. Systematic artifacts corresponding to non-cortical sources were removed by using independent component analysis (Jung et al. 2000). Condition-wise, event-related potentials (ERP) were calculated by averaging the epochs belonging to the respective condition. The central channels (F3, Fz, F4, C3, Cz, C4, P3, Pz, P4) were pooled together. The maxima/minima peak values and their timepoints (latencies) were extracted from the N100-P200 complex of the prime-induced ERPs. The minima peak values and their latencies of the negative-going deflection centered around 400 ms after target-onset (N400) were extracted from the difference waves, calculated between the two ERPs (affectively unrelated – affectively related) belonging to the same matching condition. In case of the ERPs induced by targets without preceding primes, minima values and their latencies of the negative-going deflection centered around 400 ms after target-onset were extracted from the difference waves, subtracted from each other in both directions (pleasant – unpleasant and vice versa).

#### 2.5.2 Statistical analyses

The amplitudes and the latencies as well as the mean reaction times (RT) and accuracy scores obtained from each condition by each participant were imported into the SPSS software for statistical analyses. The mean RTs as well as the accuracy scores were subjected to three-way mixed analyses of variance (ANOVA) with group (AP, NAP) as between-factor and two within-factors, namely “matching condition” with four levels (no prime, congruent, incongruent semitone, incongruent quarter-tone) and “valence” with two levels (pleasant, unpleasant). The amplitudes and latencies of the ERPs (N100, P200, N400) were subjected to two-way mixed ANOVAs with group (AP, NAP) as between-factor and “matching condition” as within-factor with three levels (congruent, incongruent semitone, incongruent quarter-tone). The amplitudes and latencies of the negative-going waves of the control difference waves were subjected to independent-samples *t*-tests. Statistical analyses were adjusted for non-sphericity using Greenhouse–Geiser Epsilon when equal variances could not be assumed. Group-interaction effects were further inspected with post-hoc analyses. The relation between group-specific ERP findings and the AP scores achieved from the pitch-labeling test were investigated using Pearson’s product-moment correlations. Correlation analyses as well as *t*-tests were run two-tailed. Effect size measures were calculated; namely the Cohen’s *d* for *t*-tests and Generalized Eta Squared (*η*^2^_G_) for ANOVAs.

## 3. Results

### 3.1 Behavioral findings

The mean reaction times (RT) and accuracy scores achieved at each condition by both samples are depicted in Figure 2. Both samples responded not only faster (*F*_1, 40_ = 63.19, *P* = 9.26×10^−10^, *η*^2^_G_ = 0.071) but also with higher accuracy (*F*_1, 40_ = 12.64, *P* = 9.87×10^−4^, *η*^2^_G_ = 0.074) to unpleasant targets than to pleasant ones. The matching condition of the primes had a group-independent influence on the RT (*F*_1.18, 47.07_ = 22.06, *P* = 9×10^−6^, *η*^2^_G_ = 0.027). The two samples responded with comparable RT, as no group effect (*F*_1, 40_ = 2.90, *P* = 0.096, *η*^2^_G_ = 0.06) or any group-interaction effects (matching x group: *F*_1.18, 47.07_ = 1.31, *P* = 0.265, *η*^2^_G_ = 0.002; valence x group: *F*_1, 40_ = 2.42, *P* = 0.128, *η*^2^_G_ = 0.003) were revealed. Regarding the accuracy scores, the two samples differed in general (group effect: *F*_1, 40_ = 4.10, *P* = 0.050, *η*^2^_G_ = 0.055) but differed also as interaction with the matching condition (*F*_1.53, 61.44_ = 3.83, *P* = 0.037, *η*^2^_G_ = 0.012) and the valence factor (*F*_1, 40_ = 4.28, *P* = 0.045, *η*^2^_G_ = 0.026). Post-hoc pairwise comparisons (Bonferroni-adjusted at the level of *α* = 0.05 for 8 paired-samples *t*-tests) of the accuracy scores between the valences at all matching conditions across both samples revealed a significant difference only in the AP sample at the incongruent semitone condition (*t*_20_ = 3.16, Bonferroni-adjusted *P* = 0.039, *d* = 0.69).

**Fig. 2.**
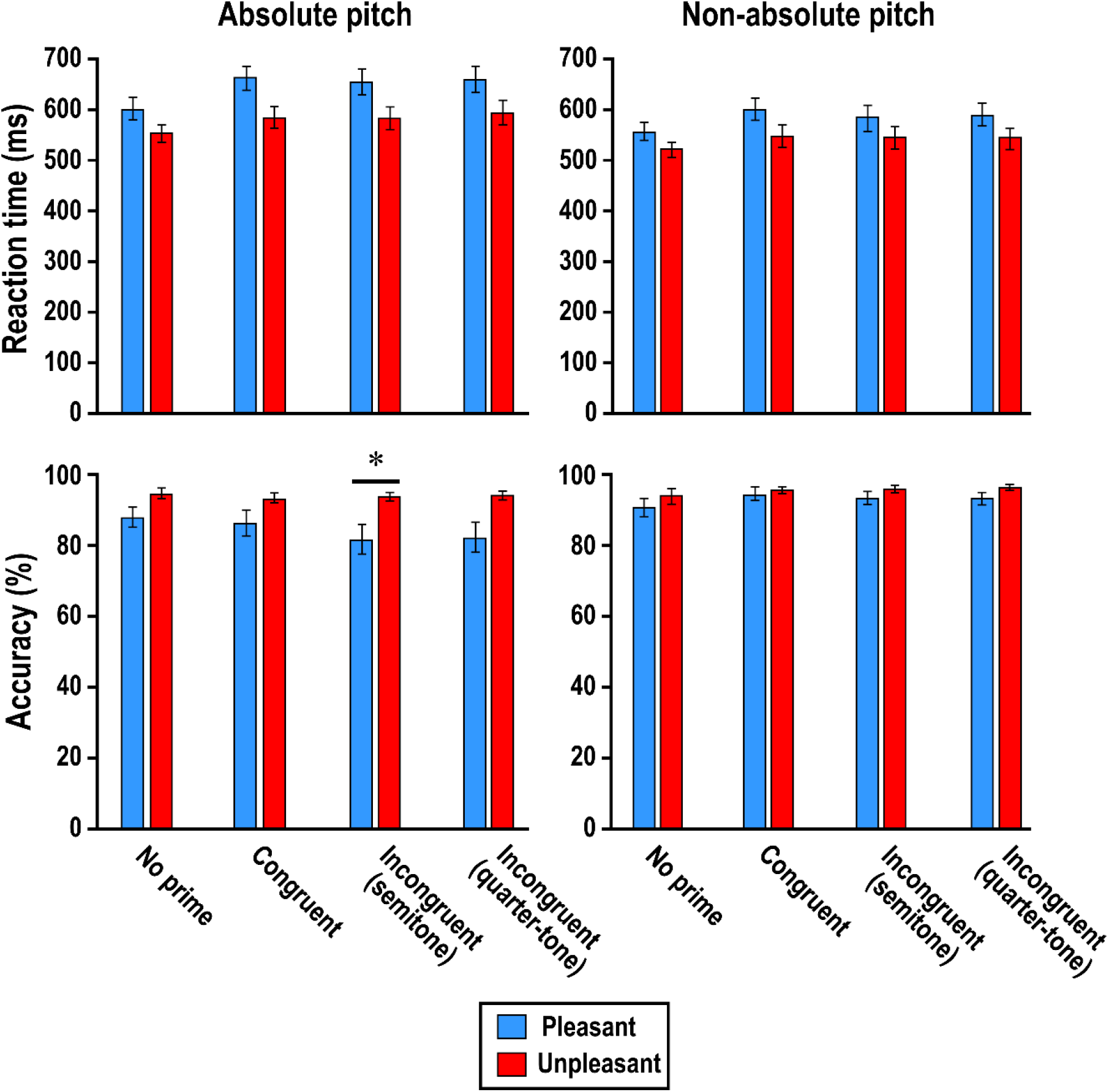
Performances achieved from the affective priming task. The mean RTs (top) as well as the accuracy scores (bottom) are depicted for each condition for the absolute pitch (left; *N* = 21) and the non-absolute pitch (right; *N* = 21) samples. The bars depict standard errors. Two-tailed Bonferroni-adjusted \**P*<0.05.

### 3.2 EEG findings

#### 3.2.1 N100-P200 complex

The ERP induced by the primes are depicted in Figure 3. and their values are plotted in Figure 4. Regarding the amplitude and the latency of the N100, no group (amplitude: *F*_1, 40_ = 0.59, *P* = 0.446, *η*^2^_G_ = 0.014; latency: *F*_1, 40_ = 0.41, *P* = 0.526, *η*^2^_G_ = 0.008), nor condition differences (amplitude: *F*_2, 80_ = 2.89, *P* = 0.061, *η*^2^_G_ = 0.004; latency: *F*_2, 80_ = 2.35, *P* = 0.102, *η*^2^ = 0.012) nor group-condition interactions (amplitude: *F* = 2.748, *P* = 0.070, *η*^2^ = 0.004; latency: *F*_2, 80_ = 0.769, *P* = 0.467, *η*^2^_G_ = 0.004) were found. Within the P200, AP possessors peaked later than NAP subjects (*F*_1, 40_ = 5.41, *P* = 0.025, *η*^2^_G_ = 0.093) but with comparable height (*F*_1, 40_ = 0.35, *P* = 0.559, *η*^2^*G* = 0.008). The P200 latencies correlated positively with the participants’ AP scores achieved from the pitch-labeling test (congruent: *r*_40_ = 0.37, *P* = 0.017; incongruent semitone: *r*_40_ = 0.31, *P* = 0.042; incongruent quarter-tone: *r*_40_ = 0.32, *P* = 0.041). The matching condition, however, had no impact on the P200 amplitudes (*F*_2, 80_ = 0.32, *P* = 0.649, *η*^2^_G_ < 0.001) nor on the P200 latencies (*F*_2, 80_ = 1.02, *P* = 0.366, *η*^2^_G_ = 0.006), and nor did it interact with the samples (amplitude: *F*_1.42, 56.63_ = 0.59, *P* = 0.531, *η*^2^_G_ = 0.002; latency: *F*_1.68, 67.35_ = 0.11, *P* = 0.862, *η*^2^ = 0.001) in this regard.

**Fig. 3.**
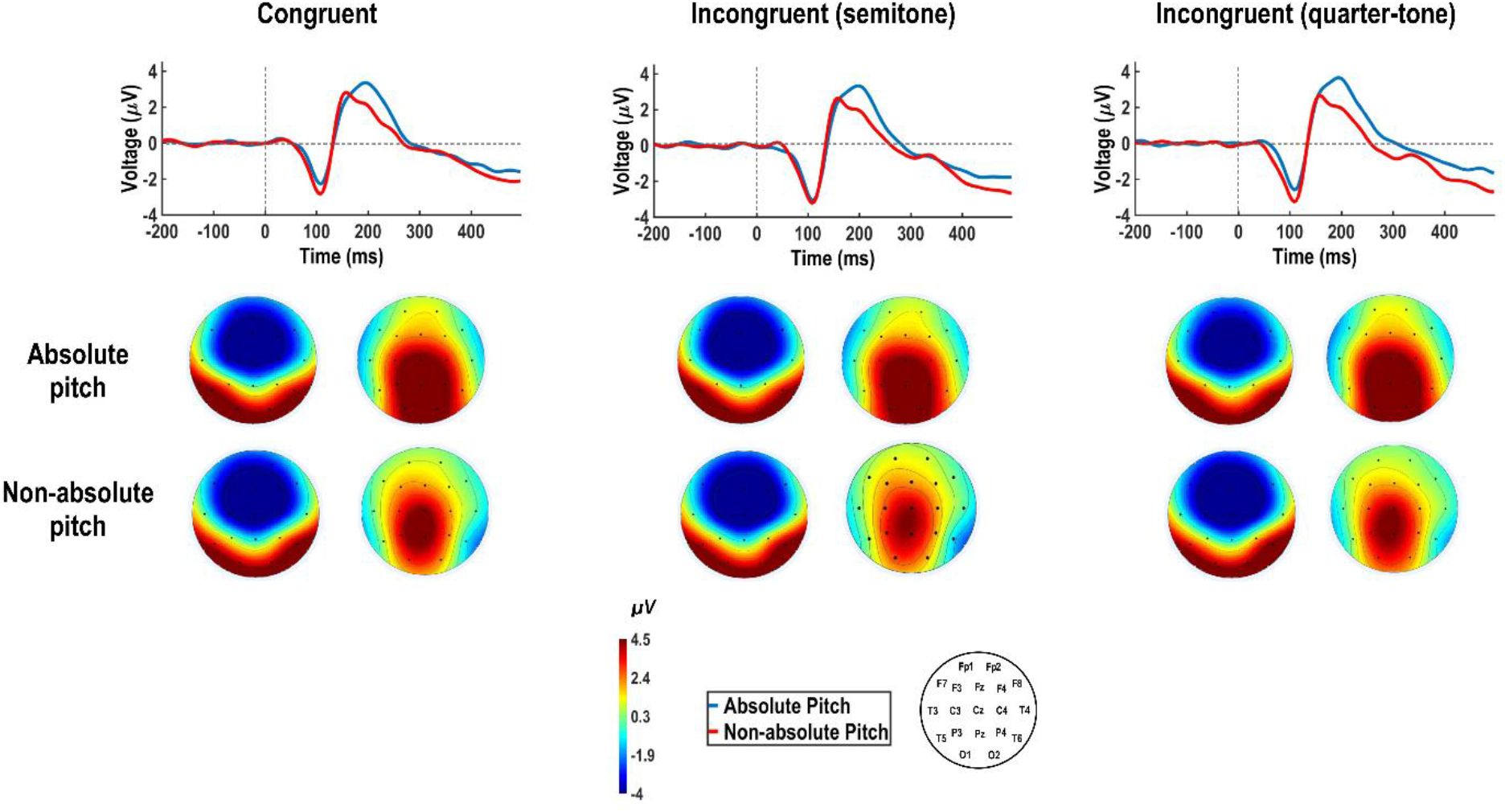
Group-averaged ERPs in response to the primes. On top, the ERPs averaged across the absolute pitch (*N* = 21) and the non-absolute pitch (*N* = 21) samples are displayed. Below, the respective current distribution of the scalps are displayed derived from the peaks of the N100 (first) and P200 (second).

**Fig. 4.**
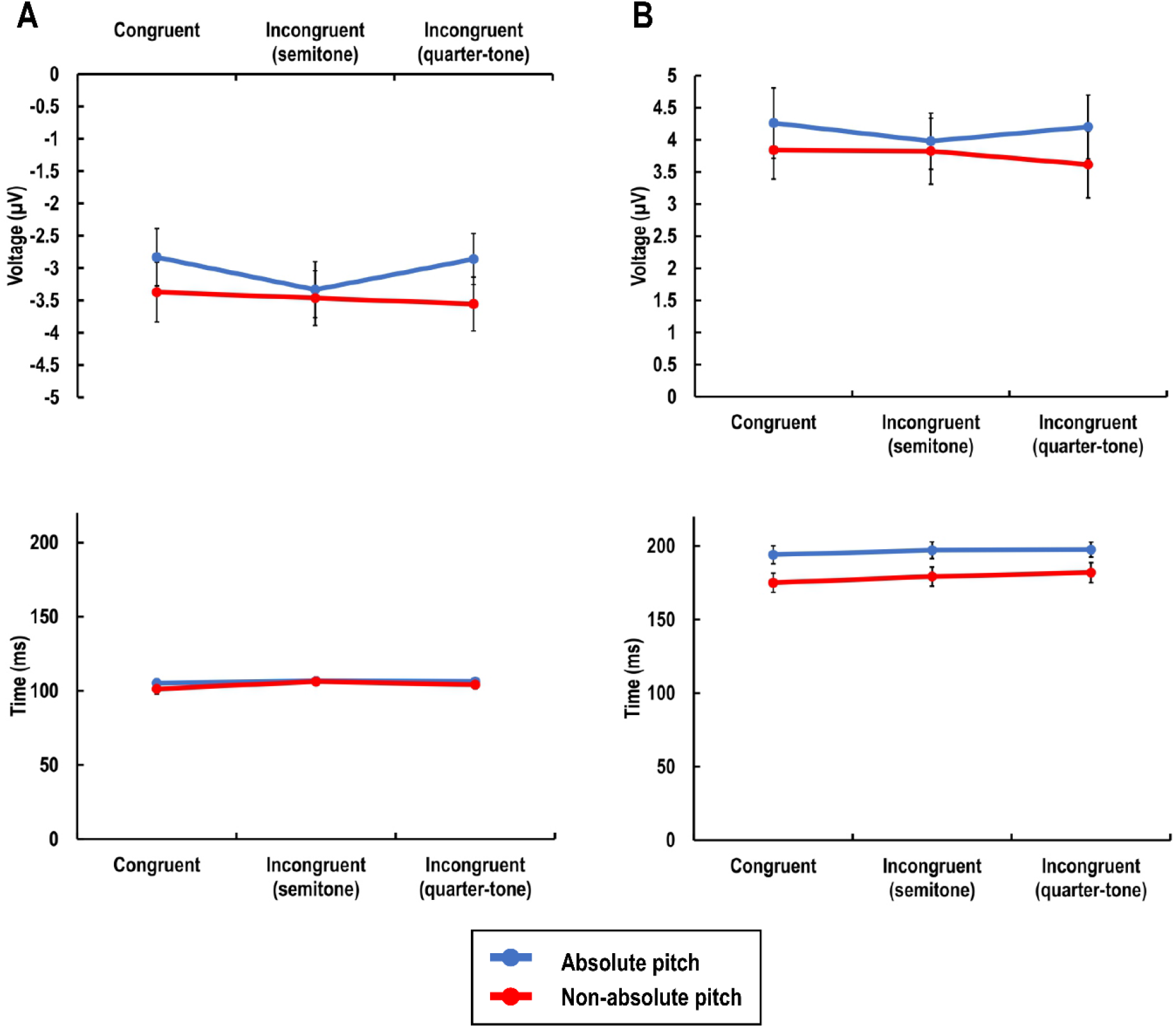
Values of the prime-induced ERPs. Plotted are the averaged amplitudes (top) and latencies (bottom) from the absolute pitch (*N* = 21) and non-absolute pitch (*N* = 21) sample across all prime matching conditions. The bars depict standard errors. A: N100. B: P200.

#### 3.2.2 N400

The ERPs induced by the targets including the difference waves calculated between the two valence factors referring to each prime matching condition are depicted in Figure 5. The values of the difference waves are plotted in Figure 6. The negative-going deflection of the difference waves calculated between the affectively related and affectively unrelated pair conditions varied as a function of the matching condition (amplitude: *F*_1.63, 65.36_ = 12.13, *P* = 1.03×10^−4^, *η*^2^_G_ = 0.148; latency: *F*_1.66, 66.43_ = 12.73, *P* = 6.3×10^−5^, *η*^2^_G_ = 0.178). No group differences were found in the amplitudes (*F*_1, 40_ = 2.30, *P* = 0.137, *η*^2^_G_ = 0.024) as well as in the latencies (*F*_1, 40_ = 1.68, *P* = 0.203, *η*^2^_G_ = 0.013). The latencies did not reveal a group-interaction effect (*F*_1.66, 66.43_ = 1.68, *P* = 0.068, *η*^2^_G_ = 0.048), whereas the amplitudes did (*F*_1.63, 65.36_ = 5.10, *P* = 0.013, *η*^2^_G_ = 0.068). The AP scores correlated negatively with the “incongruent semitone” amplitudes (*r*_40_ = – 0.35, *P* = 0.025) and trended negatively with the “incongruent quarter-tone” amplitudes (*r*_40_ = – 0.29, *P* = 0.060) but correlated positively with the “congruent” amplitudes (*r*_40_ = 0.31, *P* = 0.044). Post-hoc comparisons (Bonferroni-adjusted at the level of *α* = 0.05 for 6 paired-samples *t*-tests) of amplitudes between all matching conditions across both samples revealed significant differences only in the AP possessors between the congruent and incongruent semitone conditions (*t*_20_ = 4.85, Bonferroni-adjusted *P* = 5.88×10^−4^, *d* = 1.64) and between the congruent and incongruent quarter-tone conditions (*t*_20_ = 3.20, Bonferroni-adjusted *P* = 0.028, *d* = 1.14). Independent-samples *t*-tests across the two samples revealed a difference at the incongruent semitone condition (*t*_40_ = 2.37, uncorrected *P* = 0.023, *d* = 0.73).

**Fig. 5.**
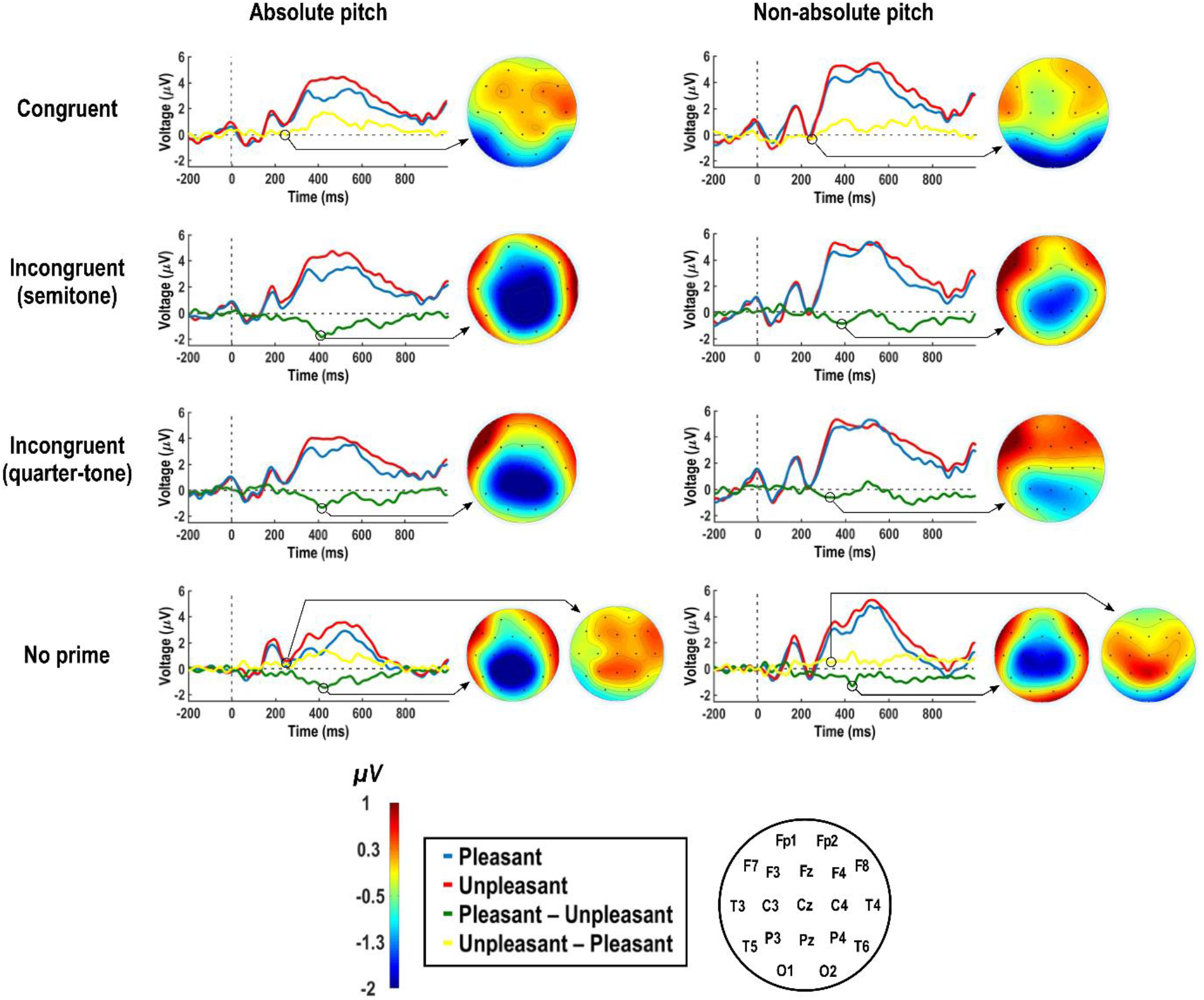
Group-averaged ERPs induced by the targets including the difference waves. The difference waves are calculated between the two ERPs (affectively unrelated – affectively related) belonging to the same matching condition. In case of the control condition (no primes), differences waves were calculated by subtracting the ERPs induced by the two targets from each other in both directions (pleasant-unpleasant and vice versa). The current distributions derived from deflection-peak of each difference curve are plotted as scalp maps. Absolute pitch (left; *N* = 21), non-absolute pitch (right; *N* = 21).

**Fig. 6.**
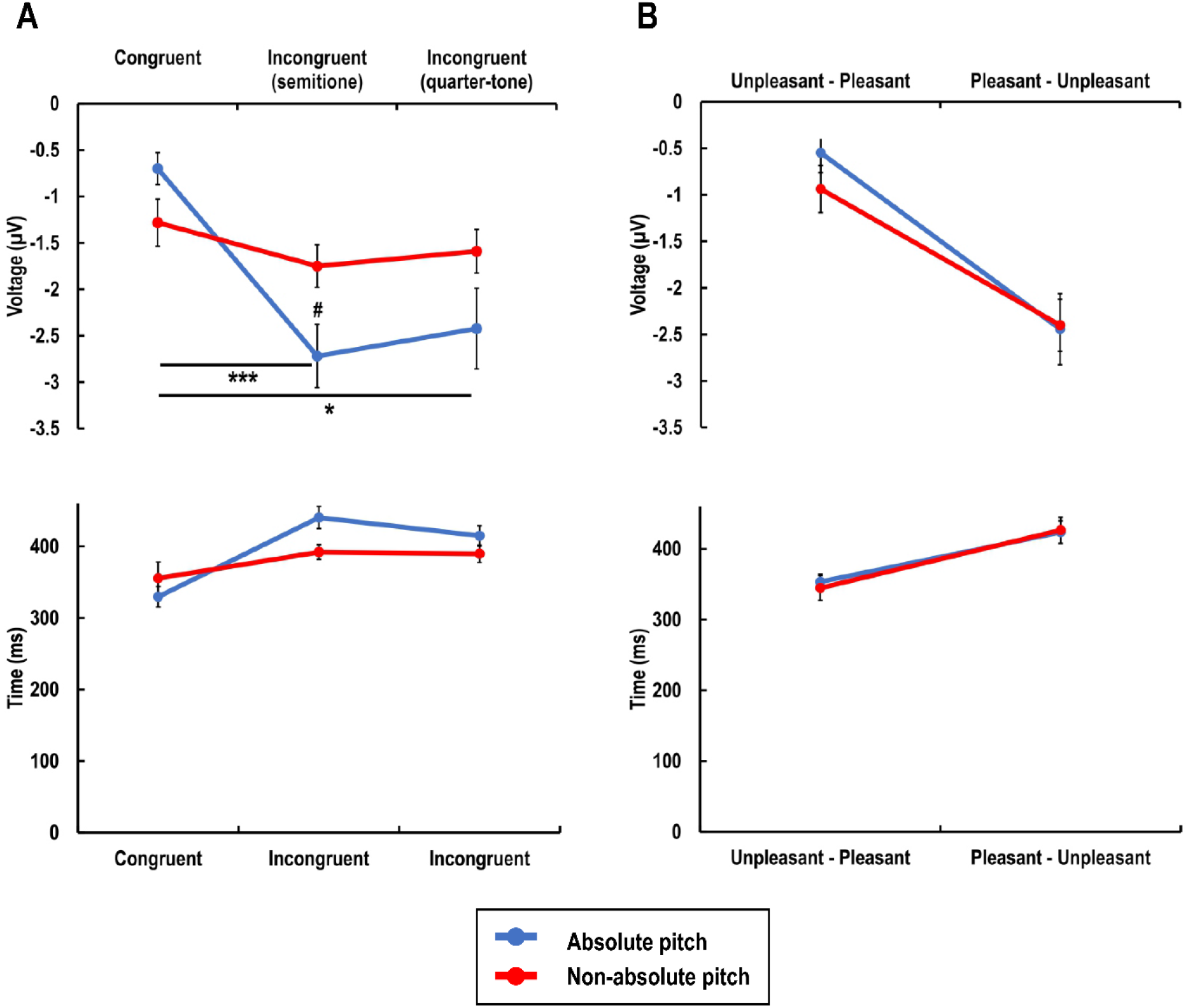
Values of the difference curves. Plotted are the group-averaged amplitudes (top) and latencies (bottom) extracted from deflection at the N400 across all prime matching (A) and control (B) conditions. Absolute pitch (*N* = 21), non-absolute pitch (*N* = 21). The bars depict standard errors. Two-tailed Bonferroni-adjusted \*\*\**P*<0.001, \**P*<0.05, and unadjusted #*P*<0.05.

#### 3.2.3 Control difference waves

Comparing the negative-going deflections of the difference wave calculated by subtraction of the ERPs induced by the unpleasant from the ERPs induced by the pleasant targets revealed no group difference in amplitude (*t*_40_ = 0.09, *P* = 0.930, *d* = 0.03) or latency (*t*_40_ = 0.14, *P* = 0.891, *d* = 0.04). Vice versa, the negative-going deflections of the difference wave calculated by subtraction of the ERPs induced by the pleasant from the ERPs induced by the unpleasant targets revealed no group difference in amplitude (*t*_40_ = 1.18, *P* = 0.244, *d* = 0.37) or latency (*t*_39.37_ = 0.36, *P* = 0.720, *d* = 0.11). The amplitudes also did not correlate with the AP scores (pleasant–unpleasant: *r*_40_ = – 0.06, *P* = 0.714; unpleasant–pleasant: *r*_40_ = 0.17, *P* = 0.294). The ERPs induced by the targets including the difference waves calculated between the two valence factors without preceding primes are also depicted in Figure 5. The values of the difference waves are plotted in Figure 6.

## 4. Discussion

The participants processed the unpleasant pictures more efficiently than the pleasant ones, as evident from the faster and more accurate responses in the unpleasant target conditions. These results mirror the well documented finding that unpleasant stimuli induce stronger affective effects than pleasant ones, supporting the idea that the processing system is sensitized to aversion (Öhman and Mineka 2001). From an evolutionary perspective, this system has adaptive value given that it favors avoidance/defense over approach response behavior. This discrepancy, referred to as negativity bias in the literature, underlies the recruitment of the subcortical route that involves the thalamus and amygdala, leading to a more rapid orienting of attention when exposed with aversive, potentially harmful events (LeDoux 1995; Ito et al. 1998; Carretié et al. 2001).

Compared to NAP subjects, the AP possessors judged the targets with accuracy scores varying as function of the prime-target pair condition. They underperformed strongest in the condition in which the pleasant targets were preceded by incongruent primes that were one semitone off. In line with the behavioral results, the N400 amplitudes differed between the two samples in dependence of the matching condition. This interaction effect was due to the conditions in which the primes were incongruent, particularly in terms one semitone. The interference in the response behavior together with the N400 effects occurring in the conditions in which the pleasant targets were preceded by incongruent primes confirm that the AP possessors processed these prime-target pairs as affectively not in line with each other, leading to the conclusion that out-of-tune musical stimuli induce aversion in AP possessors. Furthermore, these findings provide evidence that aversion is experienced more intensely when detuning reaches one semitone, possibly underlying the categorical nature of AP (Harris and Siegel 1975). Instead of relying on a sensory coding strategy, AP has shown to rather be driven by a categorical perception mechanism (Siegel 1974). Within the critical infantile period during which AP acquisition takes place, pitch-label associations are normally formed in units of semitones (Western musical system), leading to a susceptibility specified for these highly familiarized units. Thus, in comparison with within-categorical violations (e.g., quarter-tone), between-categorical violations intensify the interference. In AP possessors, detuning especially across semitone categories not only impairs music performance (Siegel 1974; Miyazaki 1993, 1995; Miyazaki and Rakowski 2002) but also, as the present findings suggest, affect the affective responses. These findings are in line with the more general relation between categorization difficulty (i.e., low fluency processing) and negative affect (Reber et al. 2004). Regarding the maturational aspect of AP, these findings are revealing given that they expand the interlinkage between AP and other developmental conditions in which AP sometimes co-exists; such as congenital synesthesia (Hänggi et al. 2008; Jäncke, Rogenmoser, et al. 2012; Gregersen et al. 2013; Bouvet et al. 2014) and blindness (Hamilton et al. 2004), autism spectrum disorder (Bonnel et al. 2003; Heaton 2003; Brenton et al. 2008; Heaton et al. 2008), Williams syndrome (Lenhoff et al. 2001). A range of auditory abnormalities are reported in Williams syndrome such as hyper- and odynacusis, auditory allodynia or attraction to certain everyday life sounds (Levitin 2005; Levitin et al. 2005). Likewise, hyperacusis and hypersensitivity to certain, especially complex, sounds have been described in autism spectrum disorder (Rosenhall et al. 1999; O’Connor 2012). Furthermore regarding common characteristics, AP possessors have shown to exhibit more autistic traits (Brown et al. 2003; Dohn et al. 2012; Wenhart, Bethlehem, et al. 2019) and rather an “autistic-typical” cognitive style, namely the processing of local information in expense of global ones (Mottron et al. 2000; Foxton et al. 2003), which is observable not only in the auditory but also in the visual domain (Costa-Giomi et al. 2001; Chin 2003; Ziv and Radin 2014; Wenhart and Altenmüller 2019; Wenhart, Hwang, et al. 2019). As another overlapping feature, imaging studies pointed out that specific hyperfunctioning, such in the cases of AP, synesthesia or other savant skills, local hyperconnectivity between the involved brain areas often underlies (Hänggi et al. 2008; Loui et al. 2010; Jäncke, Langer, et al. 2012; Mottron et al. 2013; Elmer et al. 2015; Brauchli et al. 2019). A similar phenomenon to the AP-specific aversion confirmed in this study has been described in congenital synesthesia, a condition that overlaps phenotypically and genotypically with AP (Loui et al. 2012a; Gregersen et al. 2013) and also co-occurs with autism (Mottron et al. 2013; Neufeld et al. 2013; Bouvet et al. 2014). In particular, synesthetes react aversively when exposed to inducer-concurrent pairs mismatching their familiarized idiosyncratic own (Callejas et al. 2007).

Beyond the interference reflected in the present data pattern, the AP possessors showed overall poorer accuracy scores specifically when judging pleasant pictures. A group difference in mere affective visual processing, however, was not supported by the EEG results, as the control difference curves did not differ between the two samples. Thus, the question raises to what extent AP possessors differ in affective processing not limited to out-of-tune responses or beyond the auditory dimension in general. As already mentioned, non-auditory differences between AP and NAP subjects have been reported in visual cognition, such as in the ability for spatial and binocular rivalry resolution (Costa-Giomi et al. 2001; Kim et al. 2017). Worth mentioning, a further linkage between the visual system and AP is reflected in a higher incidence rate of AP among congenitally blind (Hamilton et al. 2004). In the present study, compared to NAP subjects the AP possessors exhibited longer P200 latencies in response to the auditory-visual primes regardless of the matching condition. An ERP modulation within this relatively early component has been ascribed to the intrinsic relevance of the stimulus and to affective integration of multiple connotations (Carretié et al. 2001; Spreckelmeyer et al. 2006). Thus, the findings provide evidence that AP possessors exhibit a delay in affectively integrating musical stimuli, possible due to altered affinity to them. In earlier studies, AP possessors differed from NAP subjects in brain activation during music listening (Loui et al. 2012a) and in response to the performance of an emotional rating task during music listening (Loui et al. 2012b). In particular, AP possessors exhibited increased functional activation as well as greater functional interconnections with strongest effects around the left superior temporal gyrus including brain areas involved in multisensory integration as well as in emotion and reward processing (Loui et al. 2012b). Altogether, these findings suggest that the differences underlying the affective processing system in AP possessors are not limited to responses to out-of-tune tones.

## Appendix

The catalog numbers for the pictures used in this study are as follows.

Pleasant: 1440, 1460, 1463, 1510, 1600, 1710, 1750, 1920, 1999, 2040, 2045, 2050, 2058, 2070, 2071, 2080, 2091, 2150, 2151, 2160, 2165, 2170, 2208, 2209, 2216, 2260, 2274, 2311, 2332, 2340, 2341, 2345, 2347, 2395, 2550, 2660, 4622, 4660, 5470, 5480, 5600, 5621, 5629, 5829, 5830, 5833, 7200, 7230, 7330, 7502, 7580, 8030, 8080, 8158, 8170, 8178, 8179, 8180, 8185, 8186, 8190, 8200, 8206, 8210, 8370, 8400, 8420, 8470, 8490, 8496, 8499, 8501.

Unpleasant: 1033, 1110, 1111, 1120, 1202, 1220, 1271, 1274, 1280, 1930, 1931, 1932, 2053, 2095, 2120, 2205, 2490, 2590, 2661, 2717, 2750, 2800, 2811, 2900, 2981, 3005.1, 3015, 3030, 3064, 3071, 3100, 3140, 3160, 3181, 3220, 3225, 3230, 3300, 3301, 3350, 6020, 6021, 6520, 6555, 6563, 7380, 9006, 9007, 9008, 9040, 9042, 9043, 9075, 9102, 9140, 9182, 9183, 9265, 9301, 9320, 9325, 9373, 9405, 9490, 9530, 9561, 9570, 9571, 9582, 9584, 9592, 9594.

## Acknowledgements

This work was supported by the Swiss National Science Foundation (P1ZHP1_158642, P2ZHP1_168587, P300P1_177744 to LR and 138668, 163149 to LJ) and the National Institute on Deafness and Other Communication Disorders (RO1-DC009823 to GS).

